# NGS comparison of Downy and Powdery mildew resistance genomic regions between ‘Regent’ and ‘Red Globe’

**DOI:** 10.1101/2021.07.08.451625

**Authors:** CJ van Heerden, P Burger, JT Burger, R Prins

## Abstract

Powdery and downy mildew have a large negative impact on grape production worldwide. Quantitative trait loci (QTL) mapping projects have identified several loci for the genetic factors responsible for resistance to these pathogens. Several of these studies have focused on the cultivar ‘Regent’, which carries the resistance loci to downy mildew on chromosome 18 (*Rpv3*), as well powdery mildew on chromosome 15 (*Ren3*, *Ren9*). Several other minor resistance loci have also been identified on other chromosomes.

Here we report on the re-sequencing of the ‘Regent’ and ‘Red Globe’ (susceptible) genomes using next generation sequencing. While the genome of ‘Regent’ has more SNP variants than ‘Red Globe’, the distribution of these variants across the two genomes is not the same, nor is it uniform. The variation per gene shows that some genes have higher SNP density than others and that the number of SNPs for a given gene is not always the same for the two cultivars. In this study, we investigate the effectiveness of studying the variation of non-synonymous to synonymous SNP ratio’s between resistant and susceptible cultivars in the target QTL regions as a strategy to narrow down the number of likely candidate genes for *Rpv3*, *Ren3* and *Ren9*.

## INTRODUCTION

The wild grapevine species of America evolved separately from the domesticated European *V. vinifera* (naturally susceptible to downy and powdery mildew) and were subjected to different selective pressures, such as different pathogen exposure. Both downy and powdery mildew are endemic to the American continent and were only recently introduced to Europe. Consequently, the only downy and powdery mildew resistant *V. vinifera* cultivars (in Europe) are those derived from crosses with *non-vinifera* species, of which ‘Regent’ is a good example (Eibach & Töpfer, 2003).

Most of the R-genes identified to date contain a nucleotide-binding site (NBS) and a leucine-rich repeat (LRR) domain. These genes are effective against obligate biotrophs and hemi-biotrophic pathogens, but not necrotrophs (Glazebrook 2005). They activate a more robust defence response that often leads to localised cell death (Monaghan & Zipfel 2012). The LRR domains, involved in protein-protein interaction, show more diversity than would be expected from random genetic drift (Marone *et al*., 2013) and are under diversifying selection (Dangl & Jones 2001). It is suggested that selective pressure created by pathogens promotes the evolution of new pathogen-specificities (Marone *et al*., 2013). Specificity of resistance genes is driven by sequence variation within resistance genes (Rehmany *et al*., 2005, Dodds *et al*., 2006) and subsequently maintained in the population by selective pressure by the pathogen (Ellis *et al*., 2000). More recently, it was shown that defence-related genes evolved in parallel to the evolution of pathogens. This has been illustrated in maize for the complex rust resistance gene cluster, *Rp1* that confers race-specific resistance to *Puccinia sorghi* (Chavan *et al*., 2015). This locus has evolved to have both different copy numbers in different maize lines, as well as different sequence variants within each of these copies of the gene. Similarly, a cluster of seven resistance genes was identified on chromosome 12 of *Muscadinia rotundifolia*, a North American grapevine species. One of the genes from this cluster confers resistance to downy mildew while a second confers resistance to powdery mildew. The two genes share 86% amino acid identity (Feechan *et al*., 2013).

Given the long evolutionary distance between *V. vinifera* and the American *Vitis* species, it is likely that genes being under different selective pressures will differ from one another. Resistance to downy and powdery mildews in non-*vinifera* species evolved under selective pressure from these pathogens over an extended period and several generations. Mutations that resulted in resistance to the pathogen became fixed in the population due to selective advantage. The mutations that became fixed in the population would have altered the amino acid sequence or the expression of the gene (non-synonymous mutations). Over time, these non-synonymous mutations would accumulate as the plant evolves to combat the changing pathogen population, a process referred to as antagonistic co-evolution (Kanzaki *et al*., 2012). Synonymous mutations, on the other hand, would less likely have become fixed in the population causing a shift in the non-synonymous to synonymous SNP ratio. In contrast to *non-vinifera* species, *V. vinifera* has only recently (in evolutionary terms) been exposed to downy and powdery mildews and has not had much time to evolve resistance to these pathogens. It is therefore likely that the downy and powdery mildew resistance genes that ‘Regent’ inherited from its *non-vinifera* ancestors will have more non-synonymous variants than *V. vinifera* cultivars like ‘Red Globe’. It should however be noted that other selective pressures can lead to similar variations in the level of mutations in genes not involved in downy or powdery resistance. However, various mapping studies have identified specific genomic areas as likely locations for downy and powdery resistance genes in ‘Regent’ (Fischer *et al*., 2004; Welter *et al*., 2007; van Heerden *et al*., 2014; Zendler *et al*., 2017). These loci on chromosomes 15 and 18 differs from the chromosome 12 location reported for *Muscadinia rotundifolia*.

Based on the *non-vinifera* origin of ‘Regent’ and the possibility that its resistance could be based on a complex gene cluster, we expect that the target gene areas will contain one or more genes that have more SNPs in ‘Regent’ than in ‘Red Globe’. The difference in selection pressure during evolution would also have resulted in a difference in the ratio of non-synonymous to synonymous SNPs for some genes. This study aims to detect genes in the *Rpv3*, *Ren3* and *Ren9* loci that display an elevated number of non-synonymous variants in ‘Regent’ compared to ‘Red Globe’ (*V. vinifera*) and attempt to narrow down the number of potential resistance genes. This will aid further studies to identify the candidate resistance genes at these loci.

## MATERIALS AND METHODS

### Plant material

DNA fingerprint validated sources of ‘Regent’ and ‘Red Globe’ (van Heerden *et al*., 2014) were used for whole genome re-sequencing.

### Genome sequencing

DNA was extracted using the Plant DNA extraction kit II (Macherey Nagel, Duren, Germany; Telfer *et al*., 2013) and RNA was removed by including an RNAse1 treatment. DNA concentration was determined using the Qubit V2.

To allow for downstream data validation various sequencing platforms and chemistries (Thermo Fisher, Carlsbad, CA) were used to generate DNA sequences (Table 1). For paired-end libraries, DNA was fragmented using the Covaris S2 (Covaris, Woburn, Massachusetts, USA) prior to library construction using the 5500 SOLiD^™^ Fragment Library Core Kit (Thermo Fisher Scientific, Waltham, MA, USA). Agencourt AMPure beads (Beckman Coulter, Brea, CA, USA) were used to select DNA fragments of approximately 100 to 250 bp. Emulsion PCR and enrichment was performed on the SOLiD^™^ EZ Bead^™^ System (Thermo Fisher Scientific, Waltham, MA, USA). The paired-end protocol for sequencing 50 and 35 base reads were selected on the SOLiD^™^ 4, while the protocol for 75 and 35 base paired end reads were selected on SOLiD^™^ *5500xl*. For mate-paired sequencing, the fragmented DNA was first size selected for 1000 bp fragments using 1% agarose gel electrophoresis, followed by library construction using the 5500xl SOLiD^™^ (Thermo Fisher Scientific, Waltham, MA, USA). Emulsion PCR and enrichment were performed on the SOLiD^™^ EZ Bead^™^ System (Thermo Fisher Scientific, Waltham, MA, USA). Sequencing was performed using the mate-paired protocol for the generation of two 60 bp mated sequencing reads on the SOLiD^™^ 5500*xl*.

**TABLE 1.** Sequencing platforms and library types used for generating whole genome sequence data for ‘Regent’ and ‘Red Globe’

|  | SOLiD4 | 5500xl | 5500xl | Proton | Proton |
| --- | --- | --- | --- | --- | --- |
|  | (Paired end) | (Mate pair) | (Paired end) | (200) | (HiQ) |
| 'Regent' |  |  | X | X | X |
| 'Red Globe' | X | X |  | X | X |

For sequencing on the Ion Proton^™^, three independent simple fragment libraries were prepared using the Ion Xpress^™^ Plus Fragment Library kit for both cultivars. The libraries were templated and enriched (Ion PI^™^ Template OT2 200 Kit v2) on an Ion OneTouch^™^ 2 System prior to manual loading onto an Ion PI^™^ chip. The first two libraries for each cultivar were sequenced using the Ion PI^™^ Sequencing 200 Kit v2, while the third library for both was sequenced using the HiQ^™^ sequencing chemistry.

### Data analysis

The sequence data was quality filtered using the default parameters of the SOLiD^™^ and Ion Proton^™^ platforms respectively. No further quality trimming was performed prior to mapping of the data.

To achieve the best quality mapping results (Zhang *et al*., 2016), the Pinot Noir-derived PN40024 12× reference genome (Jaillon *et al*., 2007; http://www.genoscope.cns.fr/) was amended by adding the chloroplast (gi|91983971|ref|NC_007957.1|; Jansen *et al*., 2006) and mitochondrial (gi|224365609|ref|NC_012119.1|; Goremykin *et al*., 2009) reference genomes of grapevine using the cat command in Linux.

LifeScope^™^ V2.5 was used to analyse sequence data collected on the SOLiD^™^ platforms. All paired-end reads were mapped to the amended reference genome using LifeScope V2.5 default parameters according to the LifeScope^™^ Genomic Analysis Software 2.5 user guide. The “High stringency” setting was selected to set the variant calling parameters to values that would minimize the number of false variant calls and only SNP data was carried forward.

Torrent Suite^™^ V5.0 was used to analyse Ion Proton^™^ sequence data. For each cultivar, the three different datasets were individually mapped to the amended reference genome and the obtained binary alignment map (BAM) files combined into a single BAM file. Variant calling was performed with the variant detection plugin (tvc 5.2-16) using a modified SNP detection parameter set allowing for variant detection at a depth of cover as low as 15×, as well as allowing the use of reads with up to 20 variants per read. The variant call file (vcf) was filtered using VCFtools v0.1.14 (Danecek *et al*., 2011) to remove insertions and deletions (indels) to retain only SNP and multiple nucleotide polymorphism (MNP) data.

SNP calls were validated by comparing the SNP calls generated on the SOLiD^™^ and Ion Proton^™^ platforms for each cultivar using VCFtools v0.1.14 (Danecek *et al*., 2011). Only these validated SNPs were retained (Fig. 1) in the downstream analysis.

**Figure 1.**
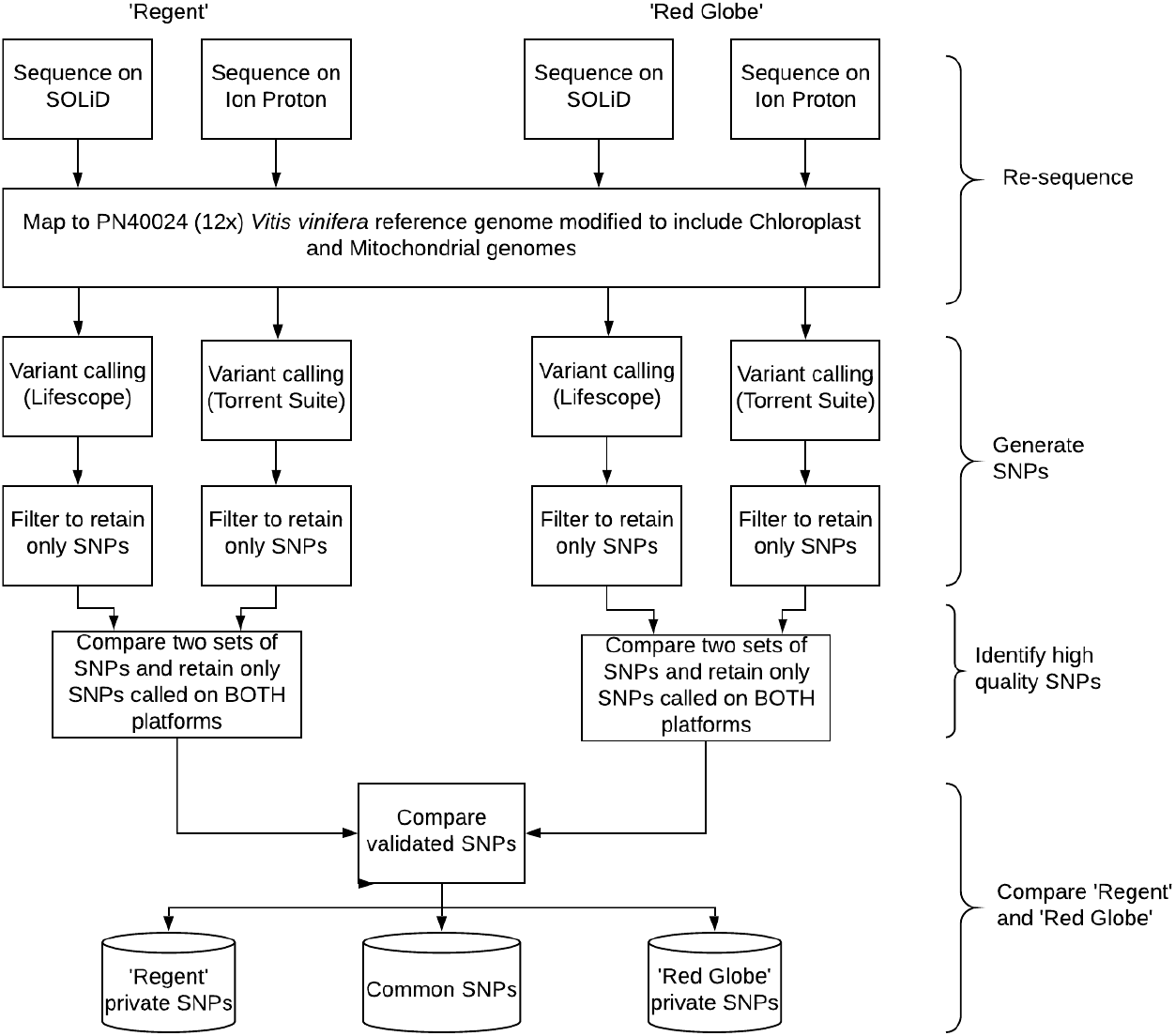
The workflow followed to produce three variant call files (vcf). The “Regent private” file contains only the SNPs that were unique to ‘Regent’ while the “Red Globe private” file contains only the SNPs that were unique to ‘Red Globe’. The “Common” file contains the SNPs found in both these cultivars. These SNPs are assumed to be private to the PN40024 12× assembled genome; positions where the PN40024 12× differs from the ancestral genome.

Subsequently, the validated SNPs for ‘Regent’ were compared to that of ‘Red Globe’ yielding three sets of SNPs. One set contains variants detected only in ‘Regent’ (‘Regent’ private), a second set contains variants detected only in ‘Red Globe’ (‘Red Globe’ private) and a third set contains variants common to both ‘Regent’ and ‘Red Globe’ (common SNPs). VCFtools v0.1.14 (Danecek *et al*., 2011) was used to calculate the private SNP density for each cultivar as well as the SNP density of the common SNPs per 1 Mbp window. The SNP densities of ‘Regent’ and ‘Red Globe’ were compared for all windows to determine if some genomic regions had a higher SNP density in ‘Regent’ than in ‘Red Globe’.

The effect of the SNPs in all three sets (‘Regent’ private, ‘Red Globe’ private and common), were classified using SNPeff (Cingolani *et al*., 2012) in conjunction with the *Vitis* genome annotation (http://plants.ensembl.org/Vitis_vinifera/Info/Index). For each gene the number of non-synonymous (ns) SNPs was calculated by combining the number of SNPs classified as initiator codon variant, missense variant, splice acceptor variant, splice donor variant, splice region variant, start lost, stop gained and stop lost. The number of synonymous (si) SNPs was determined by adding the number of SNPs classified as stop retained variant (a SNP in a stop codon that results in a stop codon) and synonymous variant. To prevent a divide by zero error, SNP counts equal to zero were replaced by 0.0001. The number of non-synonymous and synonymous SNPs and the ns/si ratio for each gene was then determined per cultivar. This information was added to a file containing gene ontologies (GO) for all annotated genes in the reference genome that were collected from UnitProt (The UniProt Consortium; http://www.uniprot.org/) and Panther (Mi *et al*., 2017; http://pantherdb.org/). In order to target regions under diversifying selection due to pathogen pressure, this list was filtered to retain only genes where the ns/si ratio for ‘Regent’ is higher than one, as well as higher than the ratio obtained for ‘Red Globe’ and the common SNP set. The resulting list of genes was then ranked according to the number of non-synonymous SNPs in ‘Regent’. To limit the analysis to potential downy and powdery mildew resistance genes, we retained only genes located between position 23 389 710 and 29 123 360 on chromosome 18 (*Rpv3* locus flanked by VVIN16-cjvh and UDV108) and between 1 125 373 and 9 410 249, and on chromosome 15 (*Ren9* and *Ren3* loci flanked by Vchr15CenGen07 and UDV116, and ScORGF15-02 and ScORA7 along with the region located between these two loci) (van Heerden *et al*., 2014; Zendler *et al*., 2017). We also include the group of contigs in the reference called “l8_random” since some of these contigs could be located in the *Rpv3* locus. To narrow the list of candidate genes further, only genes with five or more non-synonymous SNPs in ‘Regent’ were selected.

## RESULTS

### Genome sequencing results

The sequence reads for each cultivar was individually mapped to the modified PN40024 12× reference genome. The genomes of ‘Regent’ and ‘Red Globe’ were sequenced to a depth of 77× and 53×, respectively on the SOLiD^™^ platform and an average depth of 83 × for ‘Regent’ and 63× for ‘Red Globe’ was obtained for Ion Proton^™^ generated data (Table 2).

**TABLE 2.** An overview of the sequencing and mapping data using a) the SOliD4 and the 5500xl. b) Shows a similar statistic for the Ion Proton sequences.

| a) | ‘Regent’ | ‘Red Globe’ |
| --- | --- | --- |
| Number of bases mapped (Gb) | 39,40 | 26,20 |
| Number of fragments passing filters | 567 229 424 | 519 160 378 |
| Tag 1 mapped | 425 561 004 (75%) | 359 198 427 (69%) |
| Tag 2 mapped | 334 560 287 (59%) | 286 333 202 (55%) |
| Tag 1 Mean Base QV | 36 | 38 |
| Tag 2 Mean Base QV | 38 | 36 |
| Tag 1 Mean Mapping QV | 27 | 23 |
| Tag 2 Mean Mapping QV | 31 | 27 |
| Mean Coverage | 77 × | 53 × |

| b) | ‘Regent’ | ‘Red Globe’ |
| --- | --- | --- |
| Number of reads generated | 227 246 488 | 171 684 051 |
| Number of reads mapping | 221 895 789 (97.6%) | 168 636 491 (98.2%) |
| Unmapped reads | 5 350 699 | 3 047 560 |
| Number of duplicated reads | 58 067 071 | 47 266 544 |
| Number of bases mapped | 37 217 141 625 | 28 303 830 037 |
| Average read length | 165 | 166 |
| Mean base QV | 21.6 | 22.7 |
| Mean coverage | 83.82 × | 63.75 × |
| % of genome covered at >15x | 93.7% | 88.7% |

SNP calling on the SOLiD^™^ data returned a total of 5 962 303 SNPs for ‘Regent’ and 4 414 028 for ‘Red Globe’, while 7 651 348 and 6 315 089 SNPs were identified from the Ion Proton^™^ data for the respective cultivars. For ‘Regent’, 4 579 657 (9.4 SNPs per kb, or one SNP per 106.4 bases) of these SNPs were identified from both SOLiD^™^ and Ion Proton^™^ data, while 3 305 408 (7.44 SNPs per kb, or one SNP per 147.4 bases) were identified from SOLiD^™^ and Ion Proton^™^ data for ‘Red Globe’. The SNPs called on both platforms were regarded as validated (Table 3).

**TABLE 3.** Total number of SNPs identified after comparing the sequenced genomes of ‘Regent’ and ‘Red Globe’ to the Pinot Noir-derived PN40024 12× reference genome.

|  | ‘Regent’ | ‘Red Globe’ |
| --- | --- | --- |
| SNPs on Proton | 7 651 348 | 6 315 089 |
| SNPs on SOLiD | 5 962 303 | 4 414 028 |
| SNPs on both platforms (validated SNPs) | 4 579 657 | 3 305 408 |

By comparing the validated SNPs from ‘Regent’ with those from ‘Red Globe’, a list of 1 558 461 SNPs common to both cultivars was created along with lists of 3 021 196 SNPs private to ‘Regent’ and 1 746 947 SNPs private to ‘Red Globe’ (Table 4). All three SNP lists displayed a number of multi-allelic SNPs (SNPs where both alleles present at a given position differed from the sequence in the reference genome).

**TABLE 4.**
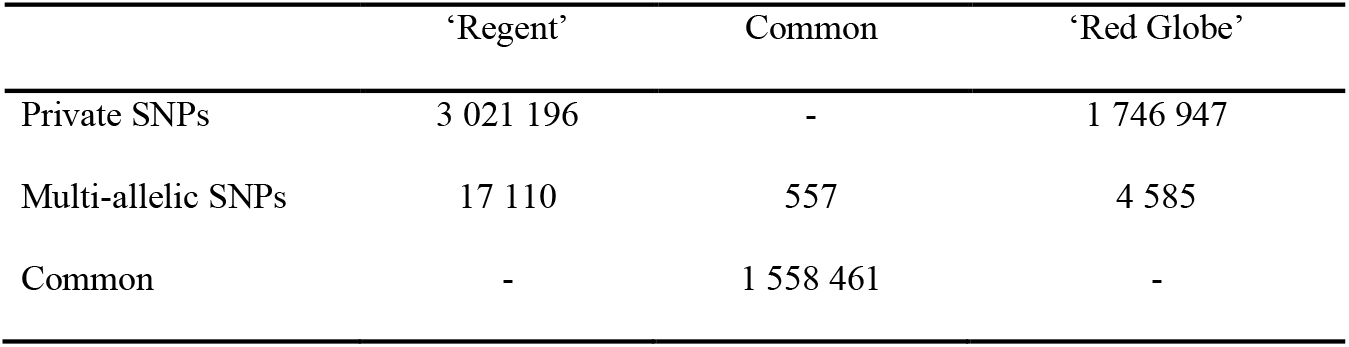
Number of validated SNPs that were found to be private to each of ‘Regent’ and ‘Red Globe’ as well as those common to both.

The SNP density of ‘Regent’, a carrier of resistance QTL from *non-vinifera* species, was compared to that of ‘Red Globe’, a pure *V. vinifera* derivative. The private SNP density of the chromosome 18 *Rpv3* region flanked by VVIN16-cjvh and UDV108 (van Heerden *et al*., 2014) in ‘Regent’ is higher than in ‘Red Globe’ (Fig. 2).

**Figure 2.**
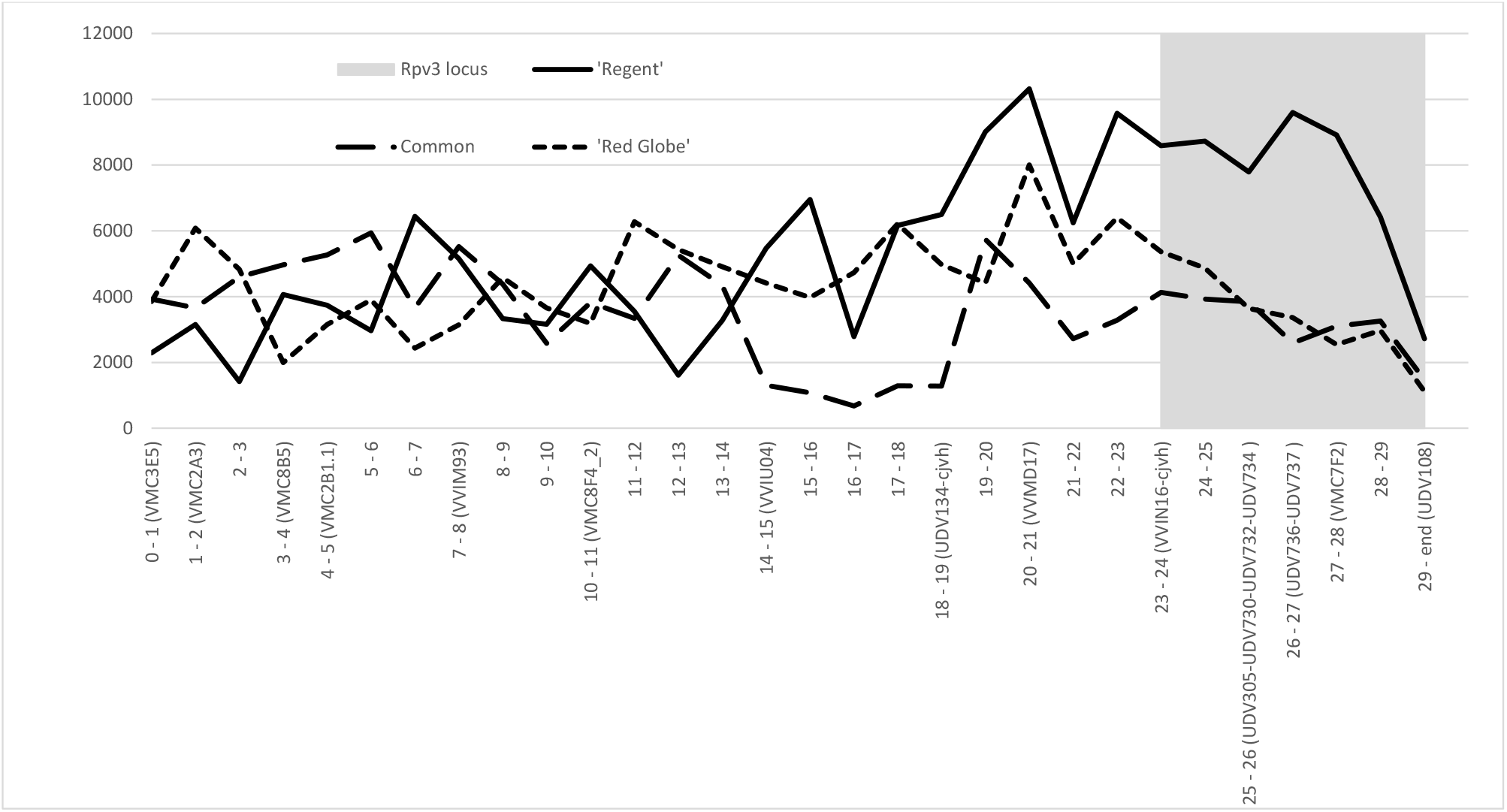
The private SNP density per megabase bin for ‘Regent’ and ‘Red Globe’ together with the density of the SNPs found to be common to both cultivars. The location of the *Rpv3* locus (grey block) and several genetic markers are indicated in relation to the bins

Annotation of the variants using SNPeff revealed that for all three sets of SNPs only 2.64 - 3.27% of the variants found were located in exons (Fig. 3) while 38.68 - 43.77% was found in intergenic regions and 13.93 - 16.32% were located in introns. The genome wide ns/si ratio was 1.09, 1.16 and 0.99 for ‘Regent’, ‘Red Globe’ and the common SNPs respectively.

**Figure 3.**
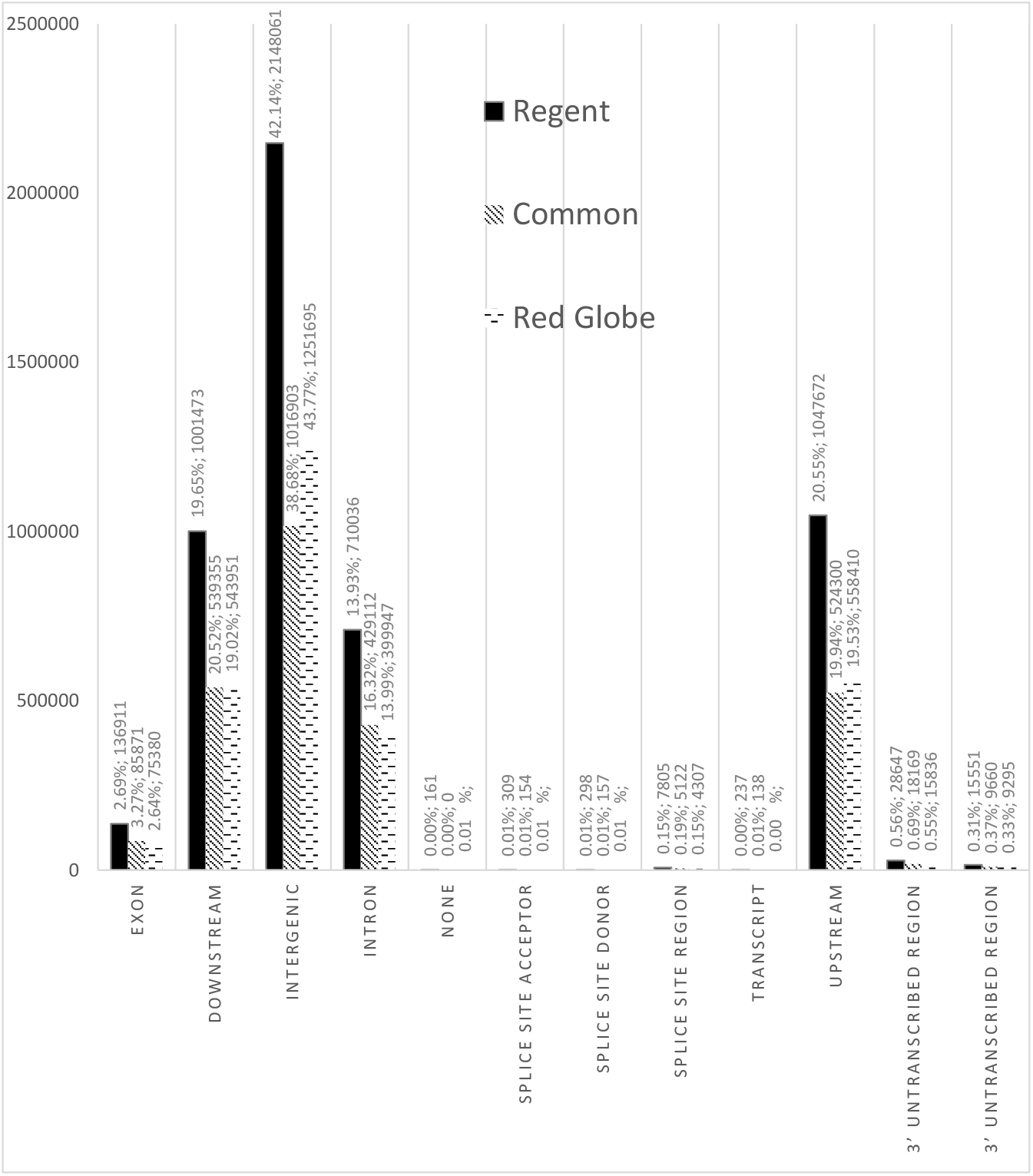
Number of private and common SNPS per type

For the remainder of the analysis, we focused on position 23 389 710 to 29 123 360 on chromosome 18 (*Rpv3* locus flanked by VVIN16-cjvh and UDV108) and position 1 125 373 to 9 410 249 on chromosome 15 (*Ren3* and *Ren9* loci flanked by Vchr15CenGen07 and UDV116, and ScORGF15-02 and ScORA7 along with the region between these two loci) (van Heerden *et al*., 2014; Zendler *et al*., 2017). We also include the group of contigs in the reference called “l8_random” since some of these contigs could be located in the *Rpv3* locus.

### *Rpv3* locus

The 350 genes located within the *Rpv3* locus revealed 3073 validated SNPs private to ‘Regent’. Of these, 1989 were non-synonymous while the remaining 1084 were synonymous (ratio 1.83). In contrast, only 1291 variants were private to ‘Red Globe’ with 827 non-synonymous and 464 synonymous (ratio 1.78). ‘Regent’ and ‘Red Globe’ had 1712 SNPs in common with 1017 nonsynonymous and 695 synonymous (ratio 1.46). A further 238 genes are located on the “chromosome” 18-random – the collection of contigs that could not be placed accurately at a specific location on chromosome 18. This collection of contigs cannot be ruled out as originating from the *Rpv3* region. Analysis of this region in ‘Regent’ revealed 995 nonsynonymous and 610 synonymous variants (ratio of 1.63) while ‘Red Globe’ had 514 nonsynonymous and 310 synonymous variants (ratio of 1.65). The set of common SNPs contained 463 nonsynonymous variants and 318 synonymous variants (ratio of 1.45).

Of the total of 588 genes located in these two chromosome 18 regions, only 154 had an ns/si ratio larger than one for ‘Regent’ while also being larger than the ns/si ratio for both ‘Red Globe’ and the set of SNPs common to both ‘Regent’ and ‘Red Globe’. Thirteen of the 588 genes had “defence response” as associated GO term, but only five of these were retained in the list of 154 genes. However, all five were in the top 10 genes when ranked according to the number of non-synonymous SNPs in ‘Regent’.

### *Ren3* and *Ren9* locus

One hundred and eighty-two genes were identified in the PN40024 12× reference genome region demarcated by the *Ren3* and *Ren9* interval markers i.e. Vchr15CenGen07 to ScORA7 on chromosome 15. These genes revealed 790 SNPs private to ‘Regent’ (502 were nonsynonymous and 288 were synonymous, an ns/si ratio of 1,74) and 514 SNPs private to ‘Red Globe’ (344 non-synonymous and 170 synonymous variants, a ratio of 1.49). ‘Regent’ and ‘Red Globe’ had only 233 SNPs in common, 143 non-synonymous and 90 synonymous (ns/si ratio of 1.58).

Only 46 of the 182 genes located in this area had an ns/si ratio larger than one for ‘Regent’ while also being larger than the ns/si ratio for both ‘Red Globe’ and the set of SNPs common to both ‘Regent’ and ‘Red Globe’.

Only nine of the 182 genes had “defence response” as GO term, but none of these genes was retained in the list of 46 genes.

## DISCUSSION

DNA extraction procedures do not only extract nuclear DNA but also extracts chloroplast and mitochondrial DNA. This DNA is then inevitably sequenced along with the nuclear DNA as illustrated by the Ion Proton^™^ dataset for ‘Red Globe’, which contained 40 821 891 (24.40%) reads that mapped to the chloroplast and mitochondrial genomes (data not shown). Preliminary read alignment comparisons using the standard ‘Pinot Noir’-derived PN40024 12× reference genome and amended genome, in which the plastid genomes are accounted for, showed that approximately 10% of the unique non-duplicate reads mapped incorrectly to the standard genome due to the absence of the chloroplast and mitochondrial genomes in the reference. Similarly, a study investigating the practicality of mining human exome sequencing data for variation in mitochondria found that mapping reads to the nuclear genome and mitochondrial genome simultaneously yielded the best result (Zhang *et al*., 2016). To improve mapping accuracy and therefore the accuracy of variant calls, we amended the nuclear PN40024 12× reference genome by concatenating it with the *Vitis vinifera* chloroplast and mitochondrial reference genomes.

Recent reports have shown that the results from different SNP calling programs can differ dramatically in the SNPs and indels detected (O’Rawe *et al*., 2013). There is a much higher concordance between SNP callers when the same mapping algorithm was used for all programs (Liu *et al*., 2013). In this study, the use of two different sequencing technologies, two different mapping programs, as well as two different variant calling programs, generated a genome wide set of validated variants for each cultivar. It is recognised that the stringent approach of retaining only SNPs identified by both workflows could lead to discarding true variants. However, given that the SNP density for ‘Regent’ (9.4 SNPs per kb) and ‘Red Globe’ (7.4 SNPs per kb) are similar to the estimated 10 SNPs per kb previously reported for *V. vinifera* (Velasco *et al*., 2007), the number of true variants discarded is likely to be very low.

Since the single read format of Ion Torrent data is not capable of detecting large indels we excluded indels from this analysis.

A total of 1 558 461 validated SNPs were common between ‘Regent’ and ‘Red Globe’ and probably represent SNPs where ‘Pinot Noir’ PN40024 12× reference genome varies from the ancestral *Vitis* genome (Table 4). The observed difference noted when comparing the number of SNPs from ‘Regent’ (4 579 657) and ‘Red Globe’ (3 305 408) is as expected, based on the *non-vinifera* ancestry of ‘Regent’ (Table 3; Eibach and Töpfer 2003; Di Gaspero *et al*., 2012). This is also evident in the high number of private SNPs observed for ‘Regent’ (3 021 196). Plotting the number of private SNPs for each cultivar along with the number of common SNPs, allowed the differences in the number of variants to be visualized on a genomic scale. A comparison of the SNP density per megabase (Mb) interval reveals that for 363 of the 465 intervals, ‘Regent’ had more private SNPs than ‘Red Globe’. These areas may represent the footprint of the non-*vinifera* ancestral DNA of ‘Regent’.

SNPeff provided annotation of the validated SNPs with regard to location and potential effect. As expected, the majority of SNPs were detected in intergenic regions (Fig. 3). A genome wide overview of the ns/si ratio revealed similar ratios of 1.09, 1.16 and 0.99 for ‘Regent’, ‘Red Globe’ and the common SNPs respectively. This suggests that there is no selection for either non-synonymous or synonymous SNPs on a genome wide scale.

## Chromosome 18 *Rpv3* and 18-random region

The private SNP density of the chromosome 18 *Rpv3* region flanked by VVIN16-cjvh and UDV108 (van Heerden *et al*., 2014) in ‘Regent’ is higher than in ‘Red Globe’ (Fig. 2). Ns/si filtering allows us to narrow down the genes of interest from 588 to 60 for further analysis. This reduced the number of defence response genes (based on associated GO terms) from 13 to only five. The remaining defence response genes are discarded because the ns/si and number of non-synonymous SNPs suggest that these genes are not under diversifying selection. This list also contains gene ontologies that are not clearly linked to defence response but could be involved in resistance-like protein serine/threonine kinase, LLR and terpene synthase domains. Interestingly, this list includes terpene synthase genes, as several terpenoids with antimicrobial properties are produced in rice (*Oryza sativa*) leaves in response to infection with *Magneportha grisea*, a pathogenic fungus (Prisic *et al*., 2004). Domains for magnesium, zink, iron and flavin adenine dinucleotide binding, oxidoreductase activity, oligopeptide transport, and cell surface receptor signalling are also represented.

The *Rpv3* locus has been studied by various groups (Di Gaspero and Cipriani 2002; Peressotti *et al*., 2010; Casagrande *et al*., 2011; Zyprian *et.al*., 2016) in order to define its location and the mechanisms involved. The NBS-LLR gene homolog, rgVrip064, has been linked to downy mildew resistance (Di Gaspero and Cipriani 2002). A BLAT search (Kent 2002) of the PN40024 12× reference genome assembly (Jaillon *et al*., 2007) reveals that this sequence is currently located on the chromosome 18-random assembly. It overlaps with the location of VIT_18s0001g06110 (F6HN37) (position 4 628 767 to 4 636 005), a gene with both a LRR and a TIR domain. However, the ns/si ratio for this gene is higher in ‘Red Globe’ than ‘Regent’ and is therefore excluded as a putative candidate in this study.

Recently, Zyprian *et al*., (2016) suggested VIT_18s0041g01790 (26 685 498 to 26 687 218) as a candidate gene for *Rpv3*. They carried out QTL mapping using a GF.GA-47-42 × ‘Villard blanc’ population and showed the resistance locus originating from ‘Villard blanc’ to be associated with marker GF18-08, a redesigned VMC7F2 at position 26 896 795. They suggested VIT_18s0041g01790 as a candidate due to its close proximity to marker GF18-08 and its leucine-rich repeat structure. While this gene falls within the ‘Regent’ *Rpv3* locus, the ns/si ratio for VIT_18s0041g01790 is higher for ‘Red Globe’ than for ‘Regent’ and was therefore filtered out.

Foria et al., (2020) identified two candidate genes (*TNL1* and *TNL2*) in the *Rpv3* locus. They postulated a sequence of events starting at an ancestral single copy *TNL* which lead to the formation of the tandem gene duplicates *TNL1* and *TNL2* which corresponds to VIT_18S0041G01330 and VIT_18S0001G01340 which was followed by a duplication of *TNL2* to form *TNL2a* and *TNL2b*. They mapped the resistance to a region that contained only *TNL2a* and *TNL2b*. By using induced *Agrobacterium tumefaciens*-mediated expression of the duplicated *TNL2* pair under the control of the CaMV 35S promoter in downy mildew sensitive leaves of *V. vinifera* ‘Syrah’ they showed that expression of this gene pair resulted in a higher density of necrotic spots and decreased sporulation. However, our analysis did not suggest VIT_18S0001G01340 (*TNL2*) as a potential candidate since it displayed a higher ns/si ratio in ‘Redglobe’ than in ‘Regent’ and the overall number of non-synonymous SNPs found only in ‘Regent’ (one) is very low while the number of non-synonymous SNPs in ‘Red Globe’ (five) is higher than that of ‘Regent’. There is only one non-synonymous SNP common to both ‘Regent’ and ‘Red Globe’. These numbers suggest that this gene has not been under diversifying selection in ‘Regent’ since the divergence of the American *Vitis sp* and *V. vinifera*. The most promising gene identified in this study is VIT_18S0041G01330 (F6I3U9), which corresponds to *TNL1* in the study by Foria et al., (2020). This gene, coding for an 1184 amino acid protein, accumulated the most non-synonymous private SNPs in ‘Regent’ (29) while only one non-synonymous private SNP was detected in ‘Red Globe’. It is located (position 25 979 007 to 26 000 298) less than 1 Mb from VMC7F2 (position 26 896 989) and GF18-08 (position 26 896 795), the most significant marker associated with the *Rpv3* locus (van Heerden *et al*., 2014; Zyprian *et al*., 2016). It is flanked by UDV305 and UDV737 (positions 24 868 044 and 26 050 509), the markers that identified seven conserved haplotypes among 226 downy mildew resistant cultivars bred from North American grapevine (Di Gaspero *et al*., 2012). It has a Toll/interleukin-1 receptor homology (TIR) domain (IPR000157), an ADP binding (GO:0043531) domain, leucine-rich repeat (LRR) domains, a P-loop containing nucleoside triphosphate hydrolase and GO terms indicating a role in signal transduction. Gene expression results for *TNL1* by Foira et al. (2020) shows that this gene (VIT_18S0041G01330) is also expressed in ‘Regent’ after inoculation with *Plasmopara viticola*. Interestingly, Casagrande *et al*., (2011) showed that the pathogenesis related (PR) genes, *PR-1* and *PR-2*, are induced by the *Rpv3*-mediated response to downy mildew infection along with an increased expression of enhanced disease susceptibility-1 (*EDS1*, a gene that mediates TIR-NBS-LRR based resistance) (Hu *et al*., 2005). The downy mildew resistance gene identified in *Muscadinia rotundifolia* is also a TIR-NB-LRR gene. This supports the selection of VIT_18S0041G01330 for further investigation.

## *Ren3* and *Ren9*

Recently, the *Ren3* interval was reduced in size and an additional resistance locus, *Ren9*, was identified (Zendler *et al*., 2017) and located in the larger interval previously called *Ren3* (Fisher *et al*., 2004). To ensure that all potential resistance genes are taken into consideration, all 182 genes located in the region from *Ren3* to *Ren9* locus were included in the analysis. Only 46 of the 182 genes had an ns/si ratio larger than one for ‘Regent’, while also being larger than the ns/si ratio for both ‘Red Globe’ and the set of common SNPs. None of these genes has “defence response” as gene ontology. VIT_15S0024G00730 (D7UB85) contains an ankyrin repeat-containing domain and three transmembrane domains. The ankyrin-repeat transmembrane protein BDA1 in *Arabidopsis thaliana* was shown to constitutively activate cell death and defence responses (Yang *et al*., 2012). In rice, the ankyrin repeat containing protein, OsPIANK1, has been shown to play a role in resistance to *Magnaporthe oryzae* (Mou *et al*., 2013). Recently, Kolodziej *et al*. (2021) showed that a gene containing 12 ankyrin repeats, *Lr14a*, confers race-specific resistance to the leaf rust pathogen (*Puccinia triticina*) of common wheat. Interestingly, a gene encoding an ankyrin-like protein was also noted in the *Rpv10* Downy mildew resistance locus on chromosome 9 (Schwander *et al*., 2012). ‘Regent’ has 39 SNPs in this gene, 30 being missense mutations, while only two missense and one sense SNP is shared between ‘Regent’ and ‘Red Globe’. ‘Red Globe’ has no private SNPs for this gene. This gene has GO terms predicting protein binding and signal transduction as possible functions and is located (1 305 408 to 1 308 910) in the *Ren9* locus (position 1 125 373 to 6 351 540). VIT_15S0024G00700, a lysine-tRNA ligase gene, is the only other gene in the top 10 genes (ranked according to the number of non-synonymous SNPs in ‘Regent’) that is located in the *Ren9* locus. VIT_15S0021G00140, involved in terpenoid biosynthesis, is the only gene in the top 10 ranked genes that is located in the *Ren3* locus as reported by Zyprian *et al*. (2016) and contains 12 SNPs private to ‘Regent’ and only one synonymous SNP private to ‘Red Globe’. The region between *Ren3* and *Ren9* contains seven genes of which two have a comparatively large number of SNPs. VIT_15S0024G01760 has 28 SNPs in ‘Regent’ and none in ‘Red Globe’ or the common SNP set. In addition to an NB-ARC domain, it also includes a winged helix-turn-helix DNA-binding and a P-loop containing nucleoside triphosphate hydrolase signature. VIT_15S0045G00530 has 23 SNPs in ‘Regent’, only two in ‘Red Globe’ and five in the common SNP set. It has no predicted domains but, similar to VIT_15S0024G01760, it has a P-loop containing nucleoside triphosphate hydrolase signature.

## CONCLUSIONS

The data reported here confirm that the ‘Regent’ genome contains a very large number of SNPs, supporting the notion that the introgression of wild *Vitis* species DNA introduced a large amount of variation not available in *V. vinifera*. We also observed significant differences in the non-synonymous to synonymous mutation ratios for various genes between ‘Regent’, containing several wild American *Vitis* species in its pedigree, and ‘Red Globe’ (*V. vinifera*, Europe). These changes are most likely the result of different selective pressures on the two continents. One of these pressures is the prevalence of downy and powdery mildew pathogens on the American continent, while it was only recently introduced to Europe. The strategy of filtering genes based on the ns/si ratio and the total number of SNPs private to ‘Regent’, produced a narrowed down list of likely candidate genes for *Rpv3*, *Ren3* and *Ren9* for further investigation.

## Author Contribution Statement

CJVH – Conceptualize manuscript, experimental layout, NGS, data analysis, bulk of manuscript preparation; PB – Obtained funding and prepared project proposal, provided all material used, edited manuscript; RP and JTB – Assisted PB with project proposal and provided major inputs in manuscript preparation.

